# Novel insights into the pig gut microbiome using metagenome-assembled genomes

**DOI:** 10.1101/2022.05.19.492759

**Authors:** Devin B. Holman, Arun Kommadath, Jeffrey P. Tingley, D. Wade Abbott

**Affiliations:** Lacombe Research and Development Centre, Agriculture and Agri-Food Canada, 6000 C&E Trail, Lacombe, AB, T4L1W1, Canada; Lethbridge Research and Development Centre, Agriculture and Agri-Food Canada, 5403 1^st^ Avenue South, Lethbridge, AB, T1J 4B1, Canada; Department of Biochemistry, University of Lethbridge, Lethbridge, AB T1K 6T5, Canada

**Author notes:** Address correspondence to Devin B. Holman.

**Keywords:** metagenome-assembled genomes, antimicrobial resistance, CAZymes, swine, gut microbiome

## Abstract

Pigs are among the most numerous and intensively farmed food-producing animals in the world. The gut microbiome plays an important role in the health and performance of swine and changes rapidly after weaning. Here, fecal samples were collected from pigs at 7 different times points from 7 to 140 days of age. These swine fecal metagenomes were used to assemble 1,150 dereplicated metagenome-assembled genomes (MAGs) that were at least 90% complete and had less than 5% contamination. These MAGs represented 472 archaeal and bacterial species, and the most widely distributed MAGs were the uncultured species *Collinsella* sp002391315, *Sodaliphilus* sp004557565, and *Prevotella* sp000434975. Weaning was associated with a decrease in the relative abundance of 69 MAGs (e.g. *Escherichia coli*) and an increase in the relative abundance of 140 MAGs (e.g. *Clostridium* sp000435835, *Oliverpabstia intestinalis*). Genes encoding for the production of the short-chain fatty acids acetate, butyrate, and propionate were identified in 68.5%, 18.8%, and 8.3% of the MAGs, respectively. Carbohydrate-active enzymes associated with the degradation of arabinose oligosaccharides and mixed-linkage glucans were predicted to be most prevalent among the MAGs. Antimicrobial resistance genes were detected in 327 MAGs, including 59 MAGs with tetracycline resistance genes commonly associated with pigs such as *tet*(44), *tet*(Q), and *tet*(W). Overall, 82% of the MAGs were assigned to species that lack cultured representatives indicating that a large portion of the swine gut microbiome is still poorly characterized. The results here also demonstrate the value of MAGs in adding genomic context to gut microbiomes.

**Importance:** Many of the bacterial strains found in the mammalian gut are difficult to culture and isolate due to their various growth and nutrient requirements that are frequently unknown. Here, we assembled strain-level genomes from short metagenomic sequences, so-called metagenome-assembled genomes (MAGs), that were derived from fecal samples collected from pigs at multiple time points. The majority of these MAGs represented bacterial species that have yet to be cultured or described thus underlining the need for cultivation studies that isolate and describe novel bacterial species. The genomic context of a number of antimicrobial resistance genes commonly detected in swine was also determined. In addition, our study connected taxonomy with potential metabolic functions such as carbohydrate degradation and short-chain fatty acid production.

## Introduction

Pork production continues to increase globally (1) despite serious challenges, such as antimicrobial use and resistance (2) and infectious disease (3), that threaten its profitability and sustainability. Microbiome research has the potential to contribute solutions to some of these issues, given a better understanding of the swine gut microbiome. The pig gut microbiome, as in most mammals, provides the host with numerous benefits including protection against pathogen colonization, aiding immune system development and maturation, and production of certain vitamins and metabolites (4). The number of unique genes within the gut microbiome also greatly exceeds those encoded within the pig genome thereby providing the host with an expanded repertoire of genes that can degrade dietary substrates (5).

Cultivation of many of the microbes found in the swine gastrointestinal tract remains difficult due to their unique, and often unknown, growth requirements. Consequently this has traditionally limited study of the mammalian gut microbiome to those microbes that can be easily grown and characterized in the laboratory (6). However, this often represents only a small fraction of the microbial diversity in the gut. In recent years, metagenome-assembled genomes (MAGs) recovered from shotgun metagenomic sequences have greatly expanded the number of microbial genomes in reference databases (7-10). Although the quality of these MAGs varies, they enable researchers to connect functional potential with microbial species and strains lacking cultured representatives.

Previously, we assessed the effects of varying weaning ages on the development of the swine gut microbiome using shotgun metagenomic sequences generated from fecal samples collected from pigs throughout the swine production cycle (11). Here, we assembled those sequences and binned the assembled contigs into MAGs, retaining only those MAGs that were at least 90% complete and had less than 5% contamination. Our main objective was to characterize the functional potential represented by those MAGs, including carbohydrate-active enzymes (CAZymes) and antimicrobial resistance genes (ARGs) and to associate those functions with individual taxa. In addition, we aimed to determine if MAGs assembled in this study are representative of the pig gut microbiome in general by using metagenomic sequences from other publicly-available swine studies.

## Results

### Metagenome-assembled genomes

From 738 Gb of shotgun metagenomic sequencing data, 87,472 MAGs with greater than 90% completeness and less than 5% contamination were recovered. After dereplication at 99% ANI (average nucleotide identity), the remaining 1,150 non-redundant MAGs represented potentially unique strains that were assigned to 360 and 472 archaeal and bacterial genera and species, respectively (Supplementary Table S1). The MAGs ranged in size from 0.74 to 6.14 Mb with an average size of 2.28 Mb (SEM ± 0.02). The 358 MAGs that were not assigned to a species in the Genome Database Taxonomy (GTDB) at a 95% ANI threshold may represent potentially novel species, 32 of which also had no genus designation. When 95% ANI was used for secondary clustering, 758 dereplicated MAGs (putative species) were produced from the original 87,472 MAGs. The vast majority of the MAGs classified using GTDB were bacteria and only 10 MAGs were assigned to the archaea, all of which were identified as methanogens. In total, 19 unique phyla were represented among the MAGs (Fig. 1). The most common species designation was *Collinsella* sp002391315 (22 MAGs), followed by *Sodaliphilus* sp004557565 (19 MAGs), *Prevotella* sp000434975 (17 MAGs), and UBA3388 sp004551865 (13 MAGs). Overall, 938 MAGs were assigned to archaeal or bacterial species that lack cultured representatives.

**Figure 1.**
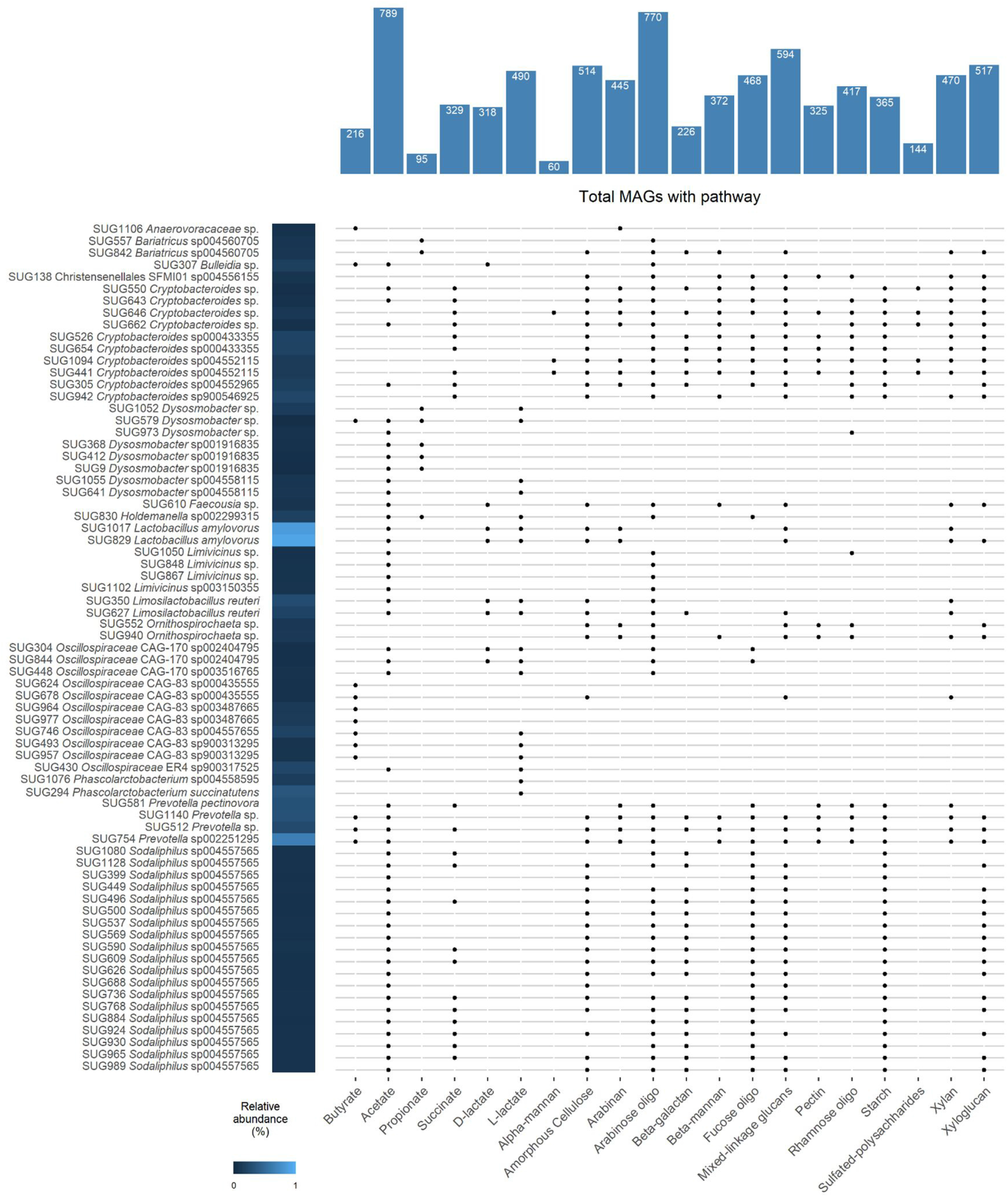
Maximum likelihood phylogenetic tree of 1,150 MAGs based on the alignment of 399 marker genes in PhyloPhlAn. MAGs colored by GTBD-tk assigned phyla are labelled in the inner ring. The outer ring indicates the number of carbohydrate-active enzymes (CAZymes) per MAG and the outer bars display the percent relative abundance (minimum = 0%; maximum = 2.73%) of each MAG in the pre- and post-weaning fecal samples.

### Functional analysis of MAGs

Functional profiling of the dereplicated MAGs using CAZymes and Kyoto Encyclopedia of Genes and Genomes (KEGG) pathways was carried out using the Distilled and Refined Annotation of Metabolism (DRAM) package. There were 6,656 unique KOs (KEGG Orthology) and 155,297 CAZymes within 281 unique CAZyme families identified among the MAGs. The average number of CAZymes per MAG was 135.0 ± 2.9 within 33.3 ± 0.5 CAZyme families (Supplementary Table S2). The glycoside hydrolases (GHs; n = 122) were most prevalent among the unique CAZyme families followed by carbohydrate-binding modules (CBMs; n = 58), glycosyltransferases (GTs; n = 55), polysaccharide lyases (PLs; n = 24), carbohydrate esterases (CEs; n = 14), and auxiliary activities (AAs; n = 8). The GH families GH2, GH3, GH13, GH23, GH25, GH31, GH32, GH36, GH73, and GH77 were most widely distributed. Many of the MAGs encoding the greatest number of CAZymes and CAZyme families belonged to the *Bacteroidaceae* family including *Bacteroides fragilis, Bacteroides heparinolyticus, Bacteroides stercoris, Bacteroides thetaiotaomicron, Bacteroides uniformis, Bacteroides xylanisolvens*, and *Phocaeicola vulgatus*. CAZymes involved in the degradation of arabinose oligosaccharides and mixed-linkage glucans were found in at least 50% of the MAGs and CAZymes predicted to be involved in the digestion of amorphous cellulose, arabinan, beta-mannan, fucose oligosaccharides, pectin, rhamnose oligosaccharides, starch, xylan, and xyloglucan were encoded by 25% or more of the MAGs (Supplementary Table S3). Mucin-degrading CAZymes were identified in only 26 MAGs, with four of these classified as *Pauljensenia hyovaginalis* and another three as *Tractidigestivibacter* sp004557505.

The production of SCFAs from carbohydrates is an important function of the gut microbiome from the perspective of the host. The most significant of these are acetate, butyrate, and propionate. Here, 68.5% of the MAGs encoded acetate-producing enzymes (acetate kinase [K00925] and phosphate acetyltransferase enzymes [K00625 or K13788]) and 8.3% had the propionate CoA-transferase gene (K01026) (Supplementary Table S3). Genes for butyrate production via the butyryl-CoA:acetate CoA-transferase (K01034, K01035) or butyrate kinase (K00634, K00929) pathways were identified in 18.8% of MAGs and included known butyrate producers, such as *Anaerostipes hadrus, Butyricimonas virosa, Butyrivibrio crossotus, Cloacibacillus porcorum, Coprococcus catus, Gemmiger formicilis, Faecalibacterium prausnitzii, Flavonifractor plautii*, and *Megasphaera elsdenii*.

Succinate is a propionate precursor that is produced by certain bacteria species. Genes encoding a fumarate reductase/succinate hydrogenase (K00239, K00240, K00241 and K00244, K00245, K00246) were identified in 28.9% of the MAGs. Known succinate producers among the MAGs with these genes included *Akkermansia muciniphila, Anaerobiospirillum succiniciproducens, Mitsuokella jalaludinii, Parabacteroides distasonis*, and *P. vulgatus*. The potential for either D-lactate or L-lactate production via lactate dehydrogenase (K00016, K03778) was detected in 55.3% of the MAGs. Genes for the production of both enantiomers of lactate were carried by 172 MAGs and over half (n = 92) of these were members of the *Treponema* genus or *Lachnospiraceae* or *Lactobacillaceae* families.

The 10 archaeal MAGs all carried the genes encoding the methyl-coenzyme M reductase complex (K00399, K00400, K00401, K00402, K03421, K03422) involved in methanogenesis (Supplementary Table S3). However, only the *Methanobacteriaceae* MAGs had the genes for the formylmethanofuran dehydrogenase complex (K00200, K00201, K00202, K00203, K00204, K00205, K11260, K11261) that is necessary for the reduction of carbon dioxide to methane. Hydrogen sulfide production in swine manure slurry has been linked to *Desulfovibrio* spp. (12) and here genes encoding the dissimilatory sulfite reductase and involved in the metabolism of sulfate were only identified in the eight *Desulfovibrionaceae* MAGs. This included *Desulfovibrio piger* and *Desulfovibrio* sp900556755 as well as one MAG classified as *Bilophila wadsworthia*. These *Desulfovibrionaceae* MAGs also carried the gene for thiosulfate reductase (K08352) which produces sulfide and sulfite through the reduction of thiosulfate.

### Antimicrobial resistance genes

The 1,150 dereplicated MAGs were screened for ARGs using the comprehensive antibiotic resistance database (CARD). A total of 327 MAGs carried at least one ARG (Supplementary Table S4), and together they accounted for 115 unique ARGs, excluding those due to point mutations. The six *Escherichia coli* MAGs contained the greatest number of ARGs (52 to 60) by a large margin. However, this is expected given that the vast majority of these ARGs are widespread within this species. ARGs conferring resistance to tetracycline (*tet* genes) are frequently among the most abundant in the gastrointestinal tract of conventionally-raised pigs and here, 59 MAGs carried at least one *tet* gene. Among the MAGs with at least one *tet* gene and an overall relative abundance of at least 0.1%, were those identified as *B. fragilis* [*tet*(Q)], *B. stercoris* [*tet*(Q)], CAG-873 sp001701165 [*tet*(Q)], *Campylobacter coli* [2 MAGs; *tet*(W/N/W), *tet*(O)], *Limosilactobacillus reuteri* [*tet*(B)], *P. vulgatus* [*tet*(Q)], *Prevotella* sp000434975 [*tet*(Q)], *Prevotella* sp000436915 [*tet* (37)], and *Streptococcus pasteurianus* [*tet*(M)].

Resistance to macrolide-lincosamide-streptogramin B (MLS_B_) antimicrobials is also often detected in the swine gut microbiome and 48 MAGs were carried at least one MLS_B_ resistance gene. Relatively abundant (≥ 0.1%) MAGs carrying one or more MLS_B_ resistance genes included *B. fragilis* [*mef*(En2)], *Catenibacterium mitsuokai* [*erm*(G)], *Clostridium* sp000435835 [*erm*(Q)], *Fusobacterium mortiferum* [*lnu*(C)], *Lactobacillus johnsonii* [2 MAGs; *erm*(B), *erm*(G)], *L. reuteri [erm*(B)*], Parabacteroides merdae* [*mef*(En2)], *P. vulgatus* [*mef*(En2)], *Treponema succinifaciens* [*erm*(F)], *Schaedlerella* sp004556565 [*lnu(C)*], SFDP01 sp004558185 [*erm*(B)], *and S. pasteurianus* [*lnu*(A), *lnu*(C), *erm*(B)].

The *vanC* cluster genes, (*vanC, vanR*_*C*_, *vanS*_*C*_, *vanT*_*C*_, *vanXY*_*C*_), which confer resistance to vancomycin were found in one MAG classified as *Enterococcus gallinarum*. In this species low-level vancomycin resistance is intrinsic due to this gene cluster (13). The beta-lactamase resistance gene *cfxA2* was identified in 13 MAGs, including six given the taxonomic designation *Sodaliphilus* sp004557565. Many of the other beta-lactamase genes detected were associated with only one bacterial species: *bla*_OXA-61_ (three *C. coli* MAGs), *bla*_TEM-1_ (one *E. coli* MAG), *cblA-1* (three *Bacteroides uniformis* MAGs), and *cepA* (one *B. fragilis* MAG). Aminoglycoside resistance genes among the relatively abundant MAGs (> 0.1%) included *aph(3’)-IIIa* in three MAGs (*C. mitsuokai, F. mortiferum*, SFDP01 sp004558185), *aac(6’)-Im* (*Blautia* spp.), *aad(6)* (SFDP01 sp004558185), *aadA* (*E. coli*), *ant(6)-Ib* (*L. johnsonii*), *aph*(2’’)-IIa (*Blautia* spp.), and *aph(6)-Id* (*E. coli*).

The location of ARGs within the MAGs was also determined to identify those ARGs co-located on the same contig as other ARGs and/or integrase/transposase sequences (Supplementary Fig. S2). The *aac(6’)-Im* and *aph(2’’)-IIa* genes were adjacent to each other in three MAGs classified as *Blautia* sp018919065, *Ruminococcus gnavus*, and CAG-238 sp. In one *T. succinifaciens* MAG, *erm*(F) and *tet*(X) were also found on the same contig as were *tet*(M) and *tet*(W/N/W) and *tet*(44) and *ant(6)-Ib* in a CAG-877 sp. MAG. In several MAGs, *lnu*(C) was co-located on the same contig as a putative transposase or integrase gene. Other ARGs potentially associated with transposases included *tet*(44) in two MAGs assigned to *Onthovivens* sp016302065 and CAG-1000 sp004552445, *tet*(M) in *Erysipelatoclostridium ramosum* and *S. pasteurianus*, and *tet*(Q) in *Onthomorpha* sp004551865 and *Prevotella* sp900548195.

### Pre-vs. post-weaning changes

As these MAGs were assembled from fecal samples taken before and after weaning it was possible to identify MAGs that were differentially abundant in the fecal microbiome of pigs immediately before weaning and 7 days post-weaning. There were 69 MAGs that were more relatively abundant in samples take just prior to weaning, the most differentially abundant of which were those classified as *Limousia pullorum, B. fragilis, E. coli, P. hyovaginalis* (*Schaalia hyovaginalis* in NCBI), and *P. vulgatus* (Supplementary Table S5). There were also six MAGs with a relative abundance greater than 0.1% in the fecal microbiomes of nursing piglets that were not detected in samples from these same pigs 7 d later. These MAGs were classified as *B. thetaiotaomicron, Bulleidia* sp., *Enterococcus faecalis, Mediterraneibacter torques, Parvimonas* sp., and *P. hyovaginalis*. Among the 140 MAGs that were most enriched in the post-weaning samples were MAGs assigned to *Copromorpha* sp., *Clostridium* sp000435835, *Fusicatenibacter saccharivorans, Intestinibacter* sp., *Oliverpabstia intestinalis, Phascolarctobacterium* sp004558595, *Prevotella* sp002251295, *Prevotella* sp004556065, and *Ruminococcus* sp003011855.

### Presence of MAGs in publicly available datasets

To determine how widely distributed the species/strains represented by the MAGs in the present study are among pigs from other studies in different geolocations, the presence and relative abundance of these MAGs within publicly available swine gut metagenomic datasets was assessed. These metagenomic sequences were from 626 fecal and cecal content samples within nine studies representing 13 different counties (Supplementary Table S6). On average, 45.5% ± 0.4% SEM of these metagenomic sequences mapped to one of the MAGs from the present study (Supplementary Table S7). Two MAGs classified as *Lactobacillus amylovorus* were the most relatively abundant overall. Other relatively abundant MAGs (>0.25%) included those identified as *B. fragilis, C. mitsuokai, L. reuteri, Phascolarctobacterium succinatutens, Prevotella pectinovora, Prevotella* sp002251295, *Prevotella* sp002300055, *Streptococcus alactolyticus*, and VUNA01 sp002299625. Metagenomic sequences from 96 MAGs were detected in 90% of these publicly available samples. Thirty-three of these MAGs were classified within the *Oscillospiraceae* family including 12 as co-abundance gene groups (CAGs), and 8 each as *Dysosmobacter* spp. and *Faecousia* spp. An additional 19 MAGs were assigned to *Sodaliphilus* sp004557565 and 8 as *Cryptobacteroides* spp.

The samples from these studies were all collected from post-weaned pigs and therefore on a non-metric multidimensional scaling (NMDS) plot of the Bray-Curtis dissimilarities the pre-weaned pig samples from the present study appear separate from the other samples (Supplementary Fig. S1). Only eight MAGs were not detected in at least one sample among all of the publicly available metagenomic samples and three of these MAGs (*Clostridium* sp., *Erysipelotrichaceae* sp., and *Negativicoccaceae* sp.,) were not identified in any of the post-weaned samples in the present study either. Overall, including the samples from the current study, there were 71 MAGs that were found in 85% of all samples (Fig. 2).

**Figure 2.**
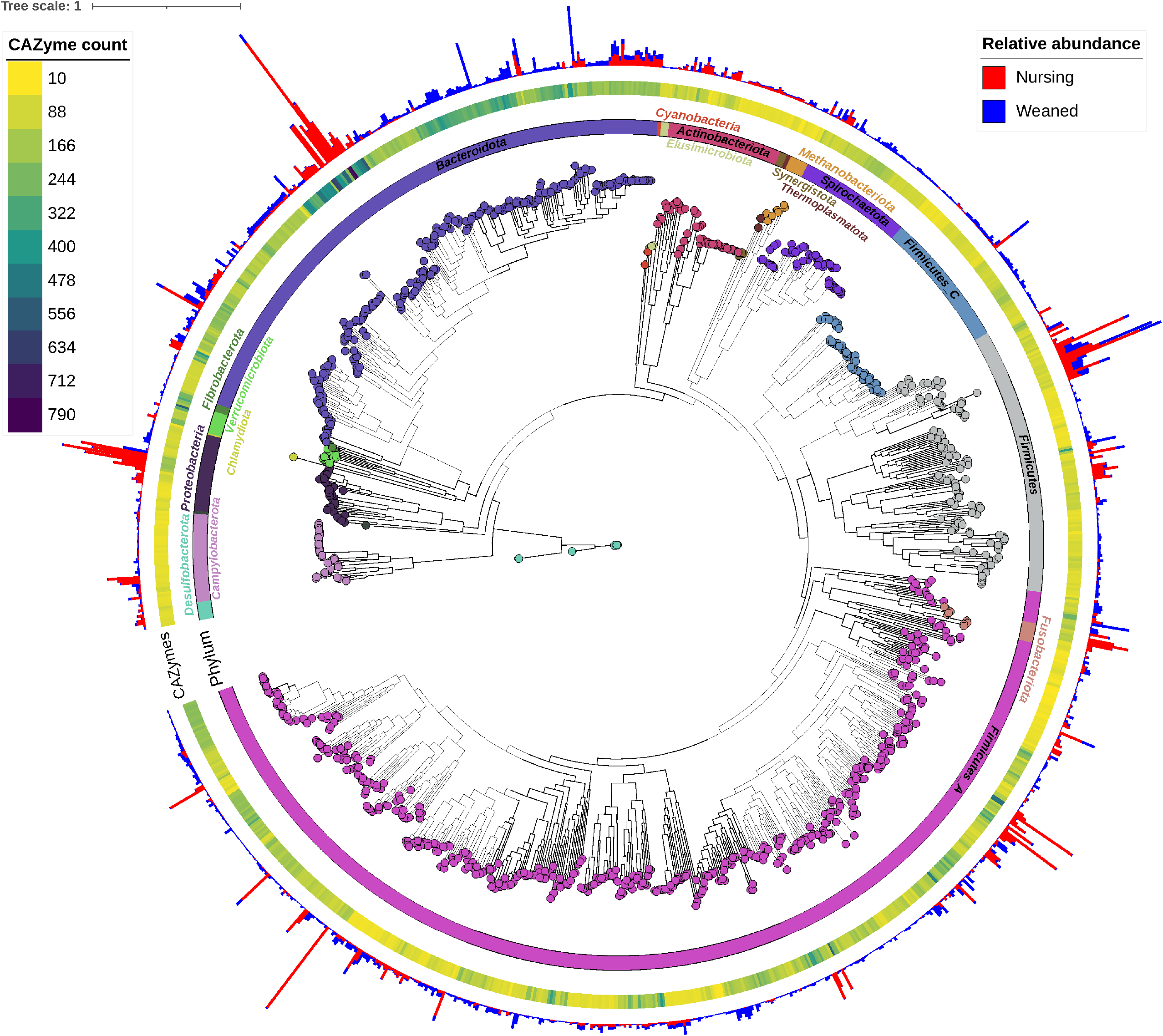
Metagenome-assembled genomes (MAGs) that were identified in 85% or more of all samples from this study and publicly available metagenome samples. The relative abundance within all of the MAGs within these samples (n = 805) is displayed as a heat map and the presence of genes encoding for pathways involved in selected short-chain fatty acid and other organic acid production, as well as polysaccharide degradation (carbohydrate-active enzymes [CAZymes]) is indicated by a dot. The total number of MAGs (n = 1,150) that encode these pathways are displayed on the top of the plot.

## Discussion

Here we assembled and analyzed 1,150 high-quality MAGs, including 358 that could not be assigned to a species and thus some of these may represent novel species. Clearly, there still exists a large fraction of the swine gut microbiome that has yet to be cultivated as demonstrated by the 938 MAGs that were not associated (>95% ANI) with a phenotypically characterized archaeal or bacterial species. Previously, we reported on changes in the pig gut microbiome in response to different weaning ages using the unassembled short reads from this study (11). We were able to recover MAGs from many of the relatively abundant species in our earlier work including *Anaeromassilibacillus senegalensis* (An172 in GTDB), *B. fragilis, E. coli, L. johnsonii, L. reuteri, P. succinatutens, P. pectinovora*, and *Subdoligranulum variabile* (*Gemmiger variabilis* in the GTDB). The exceptions were *Prevotella copri* and *Clostridioides difficile*; however, this may have been due to differences in how taxonomy was assigned to the unassembled reads vs. the MAGs as several closely related species were identified here.

After weaning, pigs are typically fed a diet that is rich in cereal grains such as corn, barley, and/or wheat which, in addition to high levels of starch, contain other polysaccharides such as cellulose, hemicellulose (e.g., mannan, mixed-linkage glucan, xylan, and xyloglucan), and pectin. These polysaccharides escape digestion by the host and are therefore available as substrates for the gut microbiome (14). As such, the pig gut microbiome carries a large repertoire of genes encoding enzymes called CAZymes that can breakdown and metabolize these polysaccharides. The CAZymes are grouped into families based on sequence similarity, although CAZymes within the same family may have different substrate specificities (15). The AAs, CEs, GHs, and PLs are the CAZyme families involved in the degradation of glycans and typically multiple CAZymes are required for the digestion of specific glycans.

Many of the MAGs encoding the greatest number of CAZymes and with the potential capacity to degrade multiple types of glycans were classified within the Bacteroidales order including *Alistipes senegalensis, Alistipes shahii, B. thetaiotaomicron, B. uniformis, B. xylanisolvens, Parabacteroides* spp., *P. vulgatus*, and *Prevotella* spp. Bacteria within this order are well documented as having a diverse and rich set of CAZymes which may be organized into groups of genes termed polysaccharide utilization loci (PUL) (16, 17). These CAZymes and PULs likely confer an advantage to these bacteria within a highly competitive ecosystem like the mammalian gastrointestinal tract. The other taxonomic group of MAGs encoding a large number of CAZymes was the *Lachnospiraceae* family that included *Acetatifactor* sp., *Blautia* sp001304935, COE1 sp., *Eisenbergiella massiliensis, Hungatella* sp005845265, and *Roseburia* sp. Some members of this family have also been reported to have gene clusters of CAZymes, regulators, and transporters that are similar to PULs (18).

The potential for the degradation of arabinan, amorphous cellulose, arabinose oligosaccharides, mixed-linkage glucans, xyloglucan, and xylan was relatively widespread among the MAGs. In pigs, diets supplemented with xylan, mixed-linkage glucans, and resistant starch have been shown to increase the relative abundance of *Blautia* spp., *Prevotella* spp. and *Lachnospiraceae* spp. (19-21). Monosaccharides produced through the action of CAZymes can then be used by the CAZyme-producer or other bacteria in the gut to generate various metabolites. In particular, the potential for short-chain fatty acid (SCFA) production through the fermentation of monosaccharides is frequently a focus of many mammalian gut microbiome studies as even in monogastric animals like pigs, up to 25% of daily energy requirements are met by SCFAs (22). Butyrate is often the SCFA of most interest as it is the primary energy source for mammalian colonic epithelial cells and can regulate apoptosis, enhance barrier function, and reduce inflammation in these cells (23, 24).

Here, 216 MAGs carried genes for butyrate production through either the butyryl-CoA:acetate CoA-transferase (*but*) or butyrate kinase (*buk*) pathways. Although several known butyrate producers are included among these MAGs such as *B. virosa, F. prausnitzii* and *M. elsdenii*, certain MAGs were assigned to bacterial species (e.g., *E. coli, E. faecalis*) that do not typically produce butyrate. Instead, these genes are likely involved in other metabolic functions or two component systems in these species. Typically, the *but* gene is more prevalent than the *buk* gene among gut bacteria (25); however, here the number of MAGs carrying either of these genes was nearly the same. There were also 17 MAGs with both *but* and *buk* genes, including *C. porcorum, F. plautii*, and *Intestinimonas massiliensis*.

Acetate and propionate, the two other physiologically important SCFAs in the mammalian gut, also have anti-inflammatory effects in addition to providing an energy source for the host (26). In the swine lower gastrointestinal tract the concentration ratio of acetate:propionate:butyrate is approximately 65:25:10 (27, 28) and here the number of MAGs (n = 788) encoding the acetate kinase and phosphate acetyltransferase genes involved in acetate production outnumbered those carrying genes for producing butyrate (n = 216) and propionate (n = 95). Bacterial species represented by these MAGs are therefore attractive targets for microbiome manipulation studies and culturing work to obtain isolates of these species for further characterization.

There were also significant shifts in the relative abundance of a large number of MAGs 7 days post-weaning. As discussed, the diet of the pigs is abruptly changed at weaning from one that is liquid and milk-based to one that is solid and based on cereal grains. This often results in a decrease in the relative abundance of *Bacteroides* and *Escherichia* spp. and an increase in the relative abundance of *Blautia, Prevotella*, and *Roseburia* spp. (29-31). Many of the differentially abundant MAGs pre- and post-weaning were assigned to these genera; however, there were also several MAGS classified as bacterial species or genera that are not known to be associated with weaning. These included MAGs enriched in post-weaning pigs that were assigned to uncultured genera or species and were also identified as potential butyrate producers such as CAG-83 sp., *Aphodosoma* sp900769035, *Copromorpha* sp., *Egerieousia* sp004561775, and UMGS1668 sp004556975. Some of these placeholder names represent bacterial taxa that have been previously reported in swine gut metagenomes and await further characterization (32, 33). One MAG classified as *E. faecalis* was relatively abundant in the nursing pig samples (0.25 ± 0.07%) but was not detected in any of the post-weaning fecal samples. *E. faecalis* was previously identified among the unassembled reads post-weaning so this MAG may represent a strain of *E. faecalis* that is unique to nursing piglets.

Binning ARGs into MAGs generated from short reads is extremely challenging as they are often flanked by repeat sequences and located on mobile genetic elements such as plasmids which have different properties (e.g., G+C content) than the chromosomal DNA of their host (34). Therefore, one can assume that ARGs identified in the MAGs here are located on the bacterial chromosome. This also explains why the number (115 ARGs) and diversity of ARGs detected in the present study was much lower than in a previous study (250 ARGs) using the same short reads that were used to assemble the MAGs here (11) as well as in the metagenome co-assembly (897 ARGs; data not shown). Despite the limitations associated with ARG binning we were able to provide genomic context for 115 ARGs including several that are relatively abundant in the swine gut such as *erm*(B), *tet*(44), *tet*(Q), and *tet*(W) (35-38).

A number of the *tet* (tetracyclines) and *erm* (MLS_B_) genes were linked to bacterial species or genera that are considered to be commensal members of the pig gut microbiome such as *Bacteroides* spp., *Clostridium* spp., *L. johnsonii, L. reuteri, Prevotella* spp., *Roseburia* spp., *Ruminococcus bromii*, and *Succinivibrio* spp. (39, 40). This may explain the extensive background level of resistance to tetracyclines and MLS_B_ antimicrobials in swine gut bacteria even in the absence of exposure to these antimicrobials, as observed here and reported in many previous studies (37, 41-43). Until relatively recently in North America, antimicrobials were often administered to all pigs in a herd for non-therapeutic purposes, namely for growth promotion (44). The gut microbiome is vertically transferred from sow to piglet and so it highly plausible that this microbiome would have been exposed to antimicrobials at some point in the past even if the pigs used in this study were not.

Several MAGs also carried ARGs conferring resistance to two or more antimicrobials. Most notable among these were a *C. coli* MAG encoding *bla*_OXA-61_, *tet*(O), and *tet*(W/N/W) and a S. *pasteurianus* MAG with *erm*(B), *lnu*(A), *lnu*(C), and *tet*(M). Both of these MAGs were also enriched in fecal samples of pre-weaned piglets. *C. coli* can be a cause of foodborne illness in humans (45) and carried by healthy pigs while *S. pasteurianus* is an opportunistic pathogen in humans and has been associated with meningitis in piglets (46). In addition, certain MAGs contained more than one ARG on the same contig, suggesting that the ARGs are linked. ARGs linked together in this manner are more likely to be co-selected and maintained within the bacterium. The aminoglycoside resistance genes *aac(6’)-Im* and *aph(2’’)-Iia* (also known as *aph(2’’)-Ib*) were adjacent to each other in three MAGs within the Clostridia class. These ARGs have previously been reported together in *Enterococcus faecium* and *E. coli* strains (47). A contig with *erm*(F) and *tet*(X) was also binned into a MAG classified as *T. succinifaciens*. These two ARGs confer resistance to macrolides and tetracyclines, respectively, and were originally described on a transposon in *B. fragilis*, although the *tet*(X) gene was reported to be inactive in this species and under anaerobic conditions (48). The *tet*(44) and *ant(6)-Ib* ARGs found here together on the same contig in a *Clostridium* sp. MAG have also been co-located on a transposon in *C. difficile* (49) and a pathogenicity island in *Campylobacter fetus* (50).

There were also a number of ARGs co-located with putative transposase or integrase genes. Transposases and integrases are enzymes that can transfer DNA segments, including ARGs, within and between bacterial genomes (51). Here, the lincosamide resistance genes *lnu*(C) and *lnu*(P) were co-located with putative transposase genes in eight different MAGs. Both *lnu*(C) and *lnu*(P) have been previously identified in *Streptococcus agalactiae* (52) and *Clostridium perfringens* (53), respectively, where they were located on the same genomic region as transposase genes. The *tet*(44), *tet*(M), *tet*(Q), and *tet*(W/N/W) genes were also detected on the same contig as putative transposase genes in certain MAGs. If these ARGs are indeed able to move between bacterial genomes it may also explain their ubiquity in swine gut metagenomes. It is possible that some of the contigs with ARGs may have been binned incorrectly given the difficulties in assembly and binning of ARGs discussed above. However, many of the ARGs were found in MAGs that were closely related to the known species range for the ARG. The use of long-read sequencing would likely increase the number of ARGs binned as well as improve the resolution of their genomic context.

We also evaluated the presence of the 1,150 MAGs from the present study within 626 swine gut metagenomes that were publicly available. Sequences aligning to 96 MAGs were identified in 90% or greater of all these samples and included 8 MAGs that were classified as *Dysosmobacter* spp. and 19 as *Sodaliphilus* sp004557565. *Dysosmobacter* is a new genus most closely related to *Oscillibacter* (54), thus explaining the absence of previous reports of this genus in the swine gut microbiome. The type species of this genus, *Dysosmobacter welbionis*, has recently been shown to partially protect against some of the negative effects of a high-fat diet when administered to mice (55). Similar to *Dysosmobacter, Sodaliphilus* is a newly described genus whose type species, *Sodaliphilus pleomorphus*, was first isolated from pig feces. Swine-derived MAGs classified as *Sodaliphilus* sp004557565 have also been recently reported (33). These results suggest that members of these genera are widespread among pigs and may represent previously unreported bacterial taxa.

## Conclusions

We recovered 1,150 high-quality MAGs from fecal metagenomes of pre- and post-weaned pigs. The MAGs described here demonstrate the vast potential of the pig gut microbiome to degrade and metabolize various glycans and of certain members to provide beneficial SCFAs to the host. In addition, the significant number of ARGs found associated with MAGs assigned to bacterial species that are typically commensals in the gut, may explain why resistance to macrolides and tetracyclines persists in the absence of antimicrobial selective pressure. The large majority of the MAGs were assigned to poorly characterized taxa and thus, there still exists a large fraction of the swine gut microbiome that has yet to be cultured. This included many bacterial species that appear to be widely disseminated among pigs from different geolocations. Future efforts focused on expanding the number of known bacterial species would greatly improve on efforts to manipulate the gut microbiome to improve production and health.

## Materials and Methods

### Experimental design

The study design and fecal sampling were previously described in Holman et al. (11). Briefly, piglets (n = 15) were assigned to be weaned at one of three ages; 14, 21, or 28 days of age. Fecal swabs were collected from the piglets at d 7, 14, 21, 28, 35, 70, and 140 days of age (n = 179). DNA was extracted using the QIAamp BiOstic Bacteremia DNA Kit (Qiagen, Mississauga, ON, Canada) and shotgun metagenomic sequencing carried out on an Illumina NovaSeq 6000 instrument (Illumina Inc., San Diego, CA, USA) with a SP flowcell (2 × 250 bp) as per Holman et al. 2021 (11).

### Bioinformatics

Metagenomic sequences were trimmed (quality score < 15 over a sliding window of 4 bp; minimum length of 50 bp) and sequencing adapters removed using Trimmomatic v. 0.38 (56). Host sequences were removed by alignment to the *Sus scrofa* genome (Sscrofa11.1) (57) using Bowtie2 v. 2.4.2-1 (58). MEGAHIT v. 1.29.0 (59) was used to co-assemble and individually assemble metagenomes. Prior to co-assembly, all metagenomic samples were normalized using BBNorm in BBTools v. 38.79 (https://sourceforge.net/projects/bbmap/). For the co-assembled metagenome, the metagenomic sequences from each sample were mapped to the co-assembly using Bowtie2 and for individual assemblies each sample was aligned to its own metagenomic assembly. These contigs in each sample with a minimum length of 2,000 bp were then binned using MetaBAT 2 (60). These bins or MAGs were assessed for quality and completeness using CheckM v. 1.1.2 (61) and those MAGs that were > 90% complete and had < 5% contamination were retained. This resulted in 2,327 MAGs from the individually assembled metagenomes and 85,145 MAGs from the co-assembled metagenomes. These MAGs were then dereplicated using dRep v. 3.2.2 (62) with primary clustering at 90% and secondary clustering at 99% ANI. These 1,150 MAGs were then used for all subsequent analyses.

Taxonomy was assigned to each MAG using GTDB-tk 2.0.0 (63) and the GTDB release 207. CoverM v. 0.6.1 (https://github.com/wwood/CoverM) (parameters: --min-read-aligned-percent 75% --min-read-percent-identity 95% --min-covered-fraction 0) was used to determine the relative abundance (coverage) of each MAG within in each metagenomic sample. A phylogenetic tree of the MAGs was constructed from the alignment of 399 marker genes in PhyloPhlAn v. 3.0.60 (64) (parameters: min_num_markers=100; f = supermatrix_aa.cfg) and visualized using iTol v6. (65). DRAM v. 1.2.4 (66) together with the KEGG (release 100, October 1, 2021) and dbCAN2 (67) databases was used to annotate the MAGs. The MAGs were also screened for ARGs using the CARD-RGI v. 5.2.0 (68). Proksee v. 1.0.0a1 (https://proksee.ca) was used to visualize the location of the ARGs within each MAG as well as potential integrases and transposases as annotated by Prokka v. 1.14.6 (69). MaAsLin2 v. 1.8.0 (70) was used to identify MAGs that were differentially abundant immediately before weaning and 7 days post-weaning. Only those MAGs with a relative abundance greater than 0.05% in these samples were included in this analysis.

Publicly available metagenomic sequences from other swine gut microbiome studies published since 2016 were downloaded and aligned to the MAGs in the present study with CoverM to assess their presence in pigs from other studies in different geographic locations. The unassembled reads as well as the MAGs from the present study are available under BioProject PRJNA629856.

## Acknowledgements

This research was supported by funding from Alberta Agriculture and Forestry grant 2018R009R and Agriculture and Agri-Food Canada.

